# External evaluation of population pharmacokinetic models for vancomycin in neonates

**DOI:** 10.1101/458125

**Authors:** Tõnis Tasa, Riste Kalamees, Jaak Vilo, Irja Lutsar, Tuuli Metsvaht

**Author notes:** **Principal investigator:** Tuuli Metsvaht. **Corresponding author:** Tõnis Tasa Address: Institute of Computer Science, University of Tartu, J. Liivi 2, 50409, Tartu, Estonia.

## Abstract

**Introduction:** Numerous vancomycin population pharmacokinetic (PK) models of neonates have been published. We aimed to comparatively evaluate a set of these models by quantifying their model-based and Bayesian concentration prediction performances using an external retrospective dataset, and estimate their attainment rates in predefined therapeutic target ranges.

**Methods:** Implementations of 12 published PK models were added in the Bayesian dose optimisation tool, DosOpt. Model based concentration predictions informed by variable number of individual concentrations were evaluated using multiple error metrics. A simulation study assessed the probabilities of target attainment (PTA) in trough concentration target ranges 10–15 mg/L and 10–20 mg/L.

**Results:** Normalized prediction distribution error analysis revealed external validation dataset discordances (global P < 0.05) with all population PK models. Inclusion of a single concentration improved both precision and accuracy. The model by Marques-Minana *et al.* (2010) attained 68% of predictions within 30% of true concentrations. Absolute percentage errors of most models were within 20-30%. Mean PTA with Zhao *et al.* (2013) was 40.4% [coefficient-of-variation (CV) 0.5%] and 62.9% (CV 0.4%) within 10–15 mg/L and 10–20 mg/L, respectively.

**Conclusion:** Predictive performances varied widely between models. Population based predictions were discordant with external validation dataset but Bayesian modelling with individual concentrations improved both precision and accuracy. Current vancomycin PK models achieve relatively low attainment of commonly recommended therapeutic target ranges.

## Introduction

For several decades, vancomycin has been one of the most widely used antibiotics in neonatal intensive care unit [1]. Therapeutic drug monitoring (TDM) is recommended due to narrow therapeutic window, especially in special patient populations like neonates, severely or critically ill etc. [2,3]. However, despite some agreement on optimal vancomycin pharmacodynamic (PD) indices and target values [4], there is little consensus on TDM based dose adjustment strategies that effectively account for high inter-individual variability [5].

The area under the curve (AUC)/ minimum inhibitory concentration (MIC) of >400 has been suggested as the best predictor of vancomycin efficacy in adults with MRSA infection [2]. However, specific software for AUC calculation as well as MIC values may not be readily available in many hospitals. Also, resulting AUC estimates are often inconsistent due to differences in choices of time-concentration curve fitting model, sampling times and estimation method [6,7]. Therapeutic considerations and relative ease of trough concentration (C_trough_) measurements make it a common proxy for therapeutic vancomycin targeting [8–10]. C_trough_ of 10 mg/L has been demonstrated as predictive and sufficient to achieve AUC>400 with high probability in paediatric populations with intermittent infusion [11–15].

A large number of population models have attempted elucidating vancomycin pharmacokinetic (PK) properties in neonates, and have applied a wide range of different structural models, covariates and methodological approaches in development [11,16–26]. These models mostly apply internal checks while only a small subset has been externally validated [27–29]. Such neglect causes prediction biases in external populations, as demonstrated by Zhao *et al.* [30]. Therefore, external assessments of model generalizability are essential for successful model portability and use in TDM applications.

Development of computational methods has introduced novel precision dosing methods to the TDM toolset [31,32]. Simple population-based empiric dose calculators can attain higher concentration prediction accuracy with Bayesian approaches that combine individual concentration measurements with population models [32,33]. Consequently, several recent software apply Bayesian methods for TDM dose optimisation including BestDose, TDMx, and DosOpt [34–36]. For example, our developed DosOpt (https://www.biit.cs.ut.ee/DosOpt) uses timed patient concentrations combined with population PK data to generate individual estimates, time-concentration curve simulations, and estimations of attainment probability for selected PK/PD target ranges.

Bayesian TDM tools require estimates of population PK that are often derived from published models. Thus, comparative model evaluations can help guide the selection of PK models for TDM tools by elucidating model prediction performance dynamics. Such evaluations that include Bayesian diagnostics, by accounting for kinetics re-adjustments with individual concentrations, have so far been performed for tacrolimus and cyclosporine but are still missing for most other drugs, including vancomycin [37,38].

We aimed to externally evaluate published neonatal vancomycin PK models to identify suitable prior population PK estimates for Bayesian vancomycin TDM. For that, we first assessed between‐ and within model predictive performances, while also considering readjustment to new patient observations. Next, we estimated vancomycin trough concentration attainment rates in therapeutic ranges to assess practical model utility.

## Methods

### Vancomycin PK models

Vancomycin PK models were included based on following criteria: (a) use of population-based, non-linear mixed effects PK modelling, (b) detailed characterisation of model structure, its compartments, and provided estimates of fixed, residual, and random effects, and (c) the average postnatal age (PNA) of the population less than 4 months.

To identify eligible models, we conducted a PubMed search by using the terms, “vancomycin AND (infant OR paediatric OR neonate) AND pharmacokinetic AND (model OR review)” until October 2017. This primary search was additionally filtered for studies published in English and involving human species. Identified articles were filtered for evidence of fitted PK models in neonatal populations. Models were further checked for availability of reported estimates, model duplications and PNA. Review articles of PK models were also identified.

### Extrenal validation dataset

The external validation dataset consisted of all patients who were admitted to the paediatric intensive care unit of Tartu University Hospital, received vancomycin, had PNA < 90 days and had at least one vancomycin concentration measurement collected for TDM between 1 January 2010 and 31 December 2015, as described previously [34]. TDM samples collected up to 2-hours before the next dose were denoted as C_trough_ and those collected 50-70 minutes after commencement of infusion were classified as peak measurements (C_peak_). Patient cases were handled as multiple individual episodes if at least a one-week gap occurred between two courses of vancomycin treatment. The following parameters were collected from hospital records: gestational age (GA), PNA, postmenstrual age (PMA), birth weight (BW), current weight (CW), serum creatinine measurement according to the Jaffe kinetic method standardized by isotope dilution mass spectrometry, use of ibuprofen or indomethacin, inotropes, and gentamicin, and respiratory support. Vancomycin was dosed according to Neofax [39] and given via a 1-hour infusion. Required dose adjustments were made at the discretion of the treating physician. Vancomycin concentrations were measured using a fluorescence polarization immunoassay (FPIA) according to the manufacturer’s instructions (Cobas Integra 400/800 Analyzer, Roche, Mannheim, Germany). The average vancomycin assay coefficient of variation was 3.5% and the lower limit of quantification was 0.74 mg/L. The Ethics Committee of the University of Tartu approved the study in Protocol No. 256/T-23. This study was a retrospective analysis with anonymized data so informed consent was not required.

### Evaluation framework

All of the external evaluation operations were performed by using Bayesian TDM software DosOpt [34]. Eligible vancomycin PK models were implemented into DosOpt and made available at https://biit.cs.ut.ee/DosOpt. Other computations were performed with R (version 3.3.3) [40].

### Population model diagnostics

We first evaluated the predictive diagnostics of population models using normalized prediction distribution error (NPDE) approach implemented in R package npde (version 2.0). A model was concordant with data when prediction errors followed a standardized Gaussian distribution. Simulation based visual predictive checks were not informative due to highly variable individually adjusted dosing schedules, sampling times and intervals [41].

We then tested if inclusion of individual concentrations improves population model fits. For accuracy, we estimated the model-wise individual mean prediction errors (MPE) as 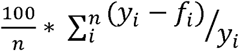. Here, *n* is number of concentrations measured for an individual, *y* is the observed and *f* the predicted concentration value. For precision, we used mean absolute prediction errors (MAPE), 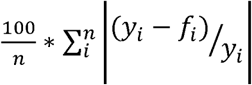. Concentration predictions were derived from the median of time-concentration curves simulated from 1000 posterior sets of PK parameters estimated with DosOpt, and were derived firstly for base population model predictions and secondly for Bayesian estimates of base models combined with full set of individual concentration measurements collected over the course of individual treatments. We also computed a priori and a posteriori adjusted-R^2^ between observed and predicted data to estimate the proportion and the magnitude of explained variability induced by inclusion of individual concentrations.

### Bayesian diagnostics

Next, we quantified predictive performances in groups of 0, 1 or 2 individual concentrations included in PK estimation. We used the median of simulated time-concentration curves to predict the next available timed concentration not included in modelling. As such, prediction accuracy was separately evaluated for individual cases with at least 1, 2 or >2 available concentrations. Accuracy for the i_th_ individual was evaluated in terms of prediction errors (PE), 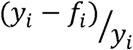, absolute errors (AE) as, |(*y_i_* – *f_i_*)|, and absolute percentage errors (APE) as 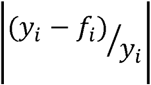. Endpoints F20 and F30 denote the percentage of individual PEs within ±20% and ±30%, respectively.

### Probability of target attainment simulations

Finally, DosOpt was used to estimate the probabilities of achieving therapeutically desired target C_trough_ ranges in a Bayesian setup given a prior PK model, individual dosing history and variable number of concentrations guiding kinetics estimation.

DosOpt optimization searched for a dose that generated time-concentration curves which attain the target C_trough_ window with highest probability. We devised an optimization scenario to find a dose for 1-hour infusion that would achieve the maximal PTA at the trough concentration, 8-hours after the start of infusion, in target C_trough_ ranges of 10–15 mg/L and 10–20 mg/L. The dose administration was simulated to start from steady-state concentration of 10 mg/L

Variance of simulated PK profiles is dependent on the variability of model between-subject parameters and residual model parameters, and the extent of shrinkage in kinetics estimation with individual data. As such, narrow targets would theoretically be better attained with models of low variability even if systematically biased. To remove that source of bias, we adjusted our concentration predictions with simulated values of expected percentage errors. First, we collected retrospectively observed mean percentage errors as described in *Population model diagnostics.* These were computed for each PK model with 0, 1 and 2 modeled concentrations that include individuals with at least 1, 2 or >2 available concentrations, respectively. Corresponding distribution parameters were estimated by fitting the values with a log-normal distribution from which 1,000 values were simulated. Beforehand, distributional assumptions were checked with Kolmogorov-Smirnov test. Therefore, simulated concentrations were scaled with model-specific log-normally distributed values that reflect expected biases in predictions. Adjustment of DosOpt concentration simulations by PE was performed according to the rearrangement of the prediction error equation: 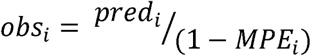, where *obs*_i_ is the adjusted concentration, *pred*_i_ is the DosOpt simulated concentration, and MPE_i_ is a simulated value from the distribution fit. The estimate for PTA is calculated as the proportion of adjusted concentration simulations as follows: 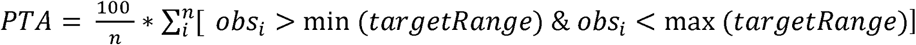, with *targetRange* being the therapeutic target range.

## Results

### External validation dataset

External validation dataset consisted of 121 patients with 149 treatment episodes. A total of 309 time-concentration points were obtained. Patient characteristics together with treatment data are presented in Table 1A and Table 1B. There was a total of 149, 84 (56.4%), and 38 (25.5%) patient treatment episodes that had at least 1, 2, and >2 vancomycin time-concentration points available, respectively. One sample was a C_peak_ (0.3% of total), 291 were denoted as C_trough_ (94% of total) and the remaining were in-between.

**Table 1.** Demographic and clinical patient data

**(A) Patient attributes (n = 149)**
| Variable (unit) | Mean (Range) | Frequencies by attribute ranges |
| --- | --- | --- |
| Postmenstrual age (weeks) | 31.7 (24.6-53.1) | <28: 38 |
|  |  | 28-33: 68 |
|  |  | 33-38: 18 |
|  |  | >38: 25 |
| Birth weight (kg) | 1.32 (0.45-4.82) | <0.8: 56 |
|  |  | 0.8-1.5: 56 |
|  |  | 1.5-2.5: 17 |
|  |  | >2.5: 20 |
| Current weight (kg) | 1.47 (0.46-5.23) | <0.8: 30 |
|  |  | 0.8-1.5: 74 |
|  |  | 1.5-2.5: 20 |
|  |  | >2.5: 25 |
| Postnatal age (days) | 19.2 (4.18-90) | <10: 55 |
|  |  | 10-25: 60 |
|  |  | 25-50: 23 |
|  |  | >50: 11 |
| Creatinine (μmol/L) | 47.3 (18-116) | <30: 35 |
|  |  | 30-50: 54 |
|  |  | 50-75: 47 |
|  |  | 75-120: 13 |

**(B)**
| Therapeutic interventions (n=149) | Inotropes | Respiratory support | Gentamicin | Ibuprofen |
| --- | --- | --- | --- | --- |
| Count (%) | 81 (54) | 132 (89) | 64 (43) | 15 (10) |

### Vancomycin PK models

Our initial search identified 96 publications. A manual review of the abstracts for these publications (performed by RK and TT) identified 24 individual studies with PK model specifications. Further filtering identified nine PK models that satisfied the inclusion criteria for original research articles (Fig. 1) [11,18–21,23–26]. Three additional models [16,17,22] were identified from review articles by Zhao *et al.* and Marsot *et al.* [30,42]. Characteristics of the models and their study populations are summarized in Electronic Supplementary Table 1.

**Fig. 1.**
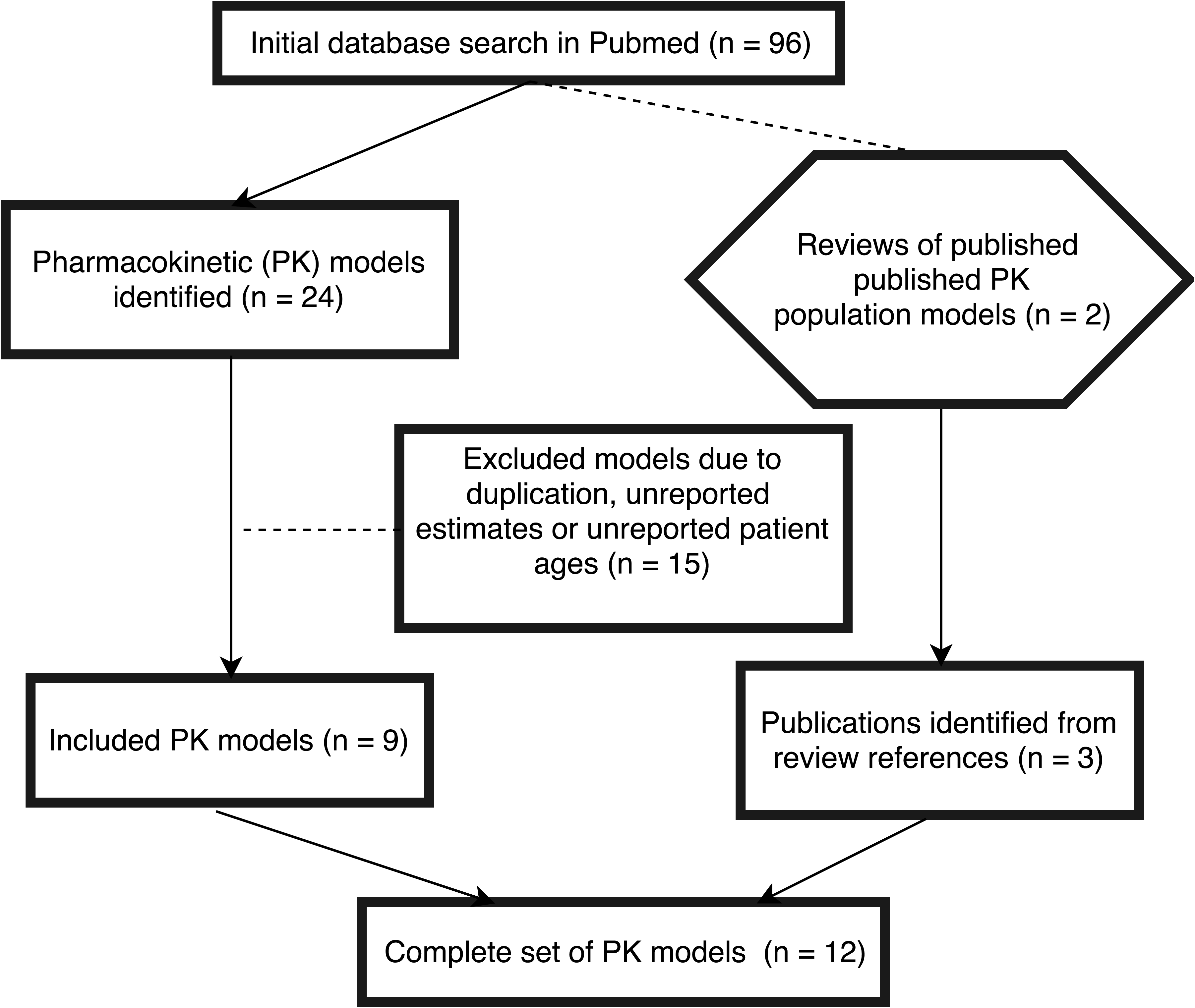
Flowchart of the model selection process.

There were several issues and ramifications regarding the models selected. First, we selected the independent vancomycin reference model and not the amikacin covariate model from De Cock *et al.* [18]. In the study by Seay *et al.* [25], no differences were reported between the 1 and 2-compartmental models. Therefore, we included the 1-compartmental model for simplicity. Since our initial simulations indicated significantly lower PTA values when a renal function component was included, the Allegaert *et al.* and Anderson *et al.* models were simplified by assuming normal renal function, i.e., a value of one [16,17]. None of the external dataset patients received spironolactone or amoxicillin-clavulcanic acid which were included as covariates in the Marques-Minana *et al.* model [21]. Patient height data were required for the Bhongsatiern *et al.* model but were not recorded in the external dataset. They were derived from Fenton growth charts based on current body weights and PNA [19,43]. These charts were also used to define SGA patients with the limit set at the 10% weight quantile for GA.

### Population model diagnostics

Population-model prediction diagnostics showed discordances of all models with our external data dataset (Fig. S1). Mean NPDE values were unbiased (Bonferroni corrected p-value threshold 0.004, n = 12) in Kimura *et al.*, De Cock *et al.*, Marques-Minana *et al.* and Zhao *et al.* models (Electronic Supplementary Table 2) [18,20,21,24]. However, we observed larger variances than expected for all models indicating worse than expected precision. Variance of Marques-Minana *et al.* NPDE values were lower than others (Fig. 2).

**Fig. 2.**
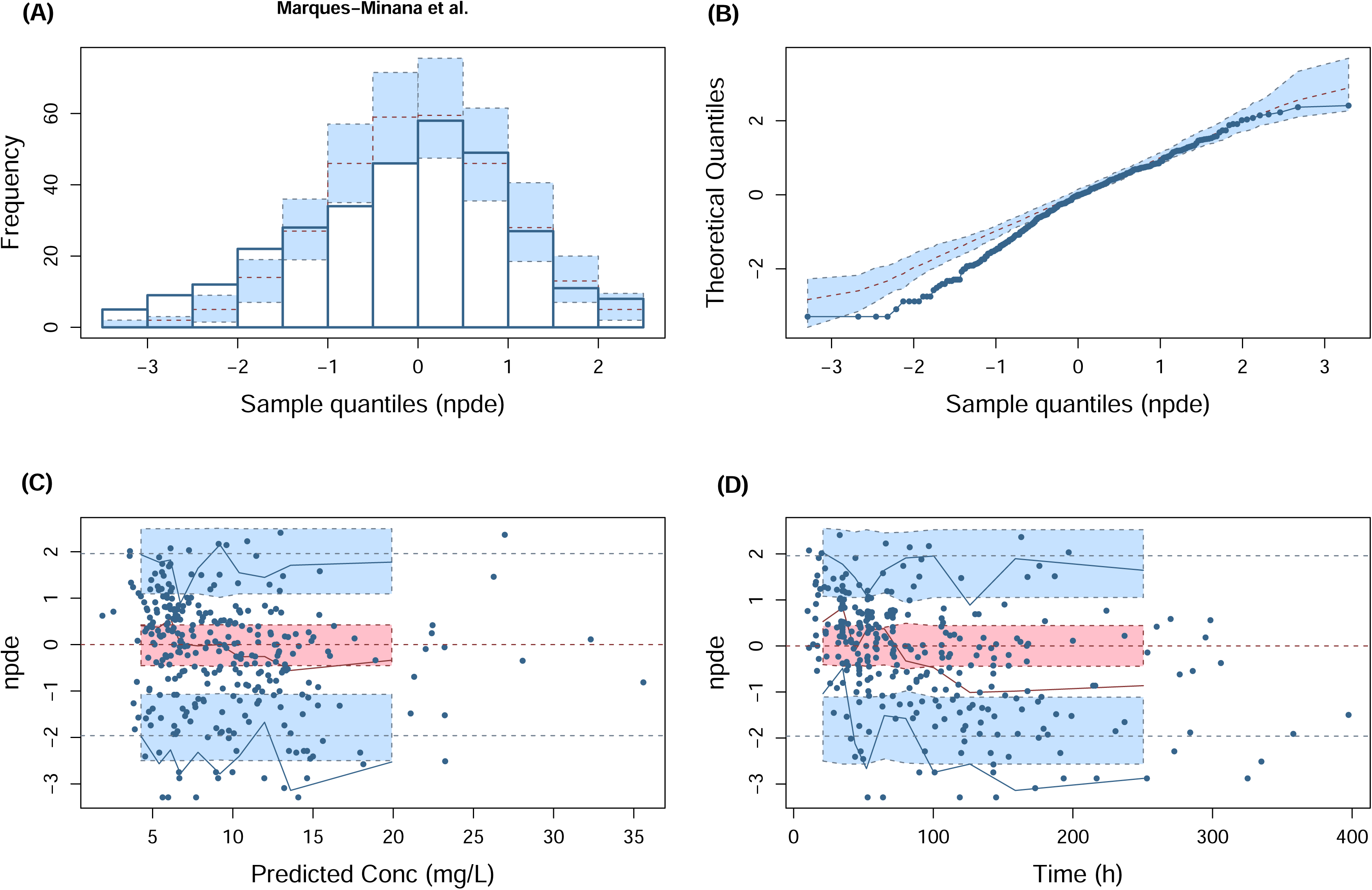
Normalized prediction distribution error (NPDE) plots of the model by Marques-Minana *et al.* (2010). (A) histogram of the distribution of the NPDE against standardized gaussian distribution (semitransparent blue fields), (B) quantile–quantile plot of the NPDE against theoretical distribution (semitransparent blue fields), (C) NPDE vs. predicted concentrations (mg/L) (D) NPDE vs. Time (h). In plot C and D, the solid red line represents the median NPDE of the observations and the semitransparent red field represents a simulation □ based 95% confidence interval (CI) for the median. Solid blue lines represent the NPDE of the observed 5th and 95th percentiles and semitransparent blue fields represent a simulation □ based 95% CI for the corresponding model predicted percentiles. The NPDE of the observations are represented by blue circles

Inclusion of individual concentration observations improved the goodness of fit of all the PK models. Modelling with individual concentrations always reduced median individual MPE values towards zero and decreased MAPE values compared to population-based predictions (Table 2). Moreover, fits of observed and predicted concentrations showed increases in the explained proportion of variability to 74% on average (Table 2, Fig. S2). This represents a 38% improvement by inclusion of all available concentrations.

**Table 2.** Mean percentage errors (MPE), mean absolute percentage errors (MAPE) and adjusted-R^2^ of the 12 population pharmacokinetics model based predictions with and without inclusion of individual concentration data. Evaluations in “All data included” group combine all individual data points with population model information, whereas “No data included” uses only population model information

| **PK model** |  |  |  |  |  |  |
| --- | --- | --- | --- | --- | --- | --- |
|  | All data included |  |  | No data included |  |  |
|  | **Median**<br>**MAPE**<br>**(SE)** | **Median**<br>**MPE**<br>**(SE)** | **R<sup>2</sup>** | **Median**<br>**MAPE**<br>**(SE)** | **Median**<br>**MPE**<br>**(SE)** | **R<sup>2</sup>** |
| Allegaert *et al.* | 0.167<br>(0.012) | 0.084<br>(0.011) | 0.753 | 0.418<br>(0.023) | 0.281<br>(0.045) | 0.296 |
| Anderson *et al.* | 0.182<br>(0.012) | 0.131<br>(0.008) | 0.779 | 0.457<br>(0.019) | 0.394<br>(0.027) | 0.413 |
| Bhongsatiern *et al.* | 0.246<br>(0.016) | 0.156<br>(0.018) | 0.72 | 0.354<br>(0.024) | 0.233<br>(0.034) | 0.508 |
| De Cock *et al.* | 0.169<br>(0.011) | 0.061<br>(0.015) | 0.606 | 0.35<br>(0.024) | 0.246<br>(0.03) | 0.236 |
| Frymoyer *et al.* | 0.159<br>(0.011) | 0.345<br>(0.022) | 0.787 | 0.35<br>(0.022) | 0.482<br>(0.028) | 0.403 |
| Grimsley and Thomson | 0.367<br>(0.018) | 0.046<br>(0.022) | 0.666 | 0.523<br>(0.021) | 0.072<br>(0.049) | 0.448 |
| Kimura *et al.* | 0.209<br>(0.016) | -0.087<br>(0.02) | 0.739 | 0.399<br>(0.3) | -0.209<br>(0.058) | 0.352 |
| Lo *et al.* | 0.173<br>(0.014) | 0.057<br>(0.008) | 0.706 | 0.396<br>(0.033) | 0.233<br>(0.044) | 0.198 |
| Marques-Minana *et al.* | 0.134<br>(0.011) | 0.07<br>(0.007) | 0.843 | 0.348<br>(0.022) | 0.324<br>(0.032) | 0.352 |
| Oudin *et al.* | 0.165<br>(0.011) | 0.014<br>(0.009) | 0.789 | 0.409<br>(0.02) | 0.191<br>(0.054) | 0.418 |
| Seay *et al.* | 0.217<br>(0.015) | -0.196<br>(0.042) | 0.705 | 0.933<br>(0.084) | -0.912<br>(0.136) | 0.295 |
| Zhao *et al.* | 0.173<br>(0.012) | 0.083<br>(0.013) | 0.776 | 0.319<br>(0.019) | 0.202<br>(0.036) | 0.428 |

### Bayesian diagnostics

Medians of APE estimated with all population models were above 31%. Population-based PEs were consistently closer to zero (t-test, P = 0.006) in all assayed patients (n = 149) compared to its subgroup with >2 concentration measurements (n = 38). Precision estimates improved when at least a single concentration (n = 84) was included in estimation, as demonstrated by decreased APE values (t-test, P = 5E-4) below 25% for seven models. Accuracy also improved as concentration inclusion approximated all models’ PE values towards zero (Fig. 3). Inclusion of a second concentration (n = 38) did not induce statistically significant changes to PEs or APEs, regardless of nominal improvement. Results of Bayesian diagnostics are reported in Electronic Supplementary Table 3.

**Fig. 3.**
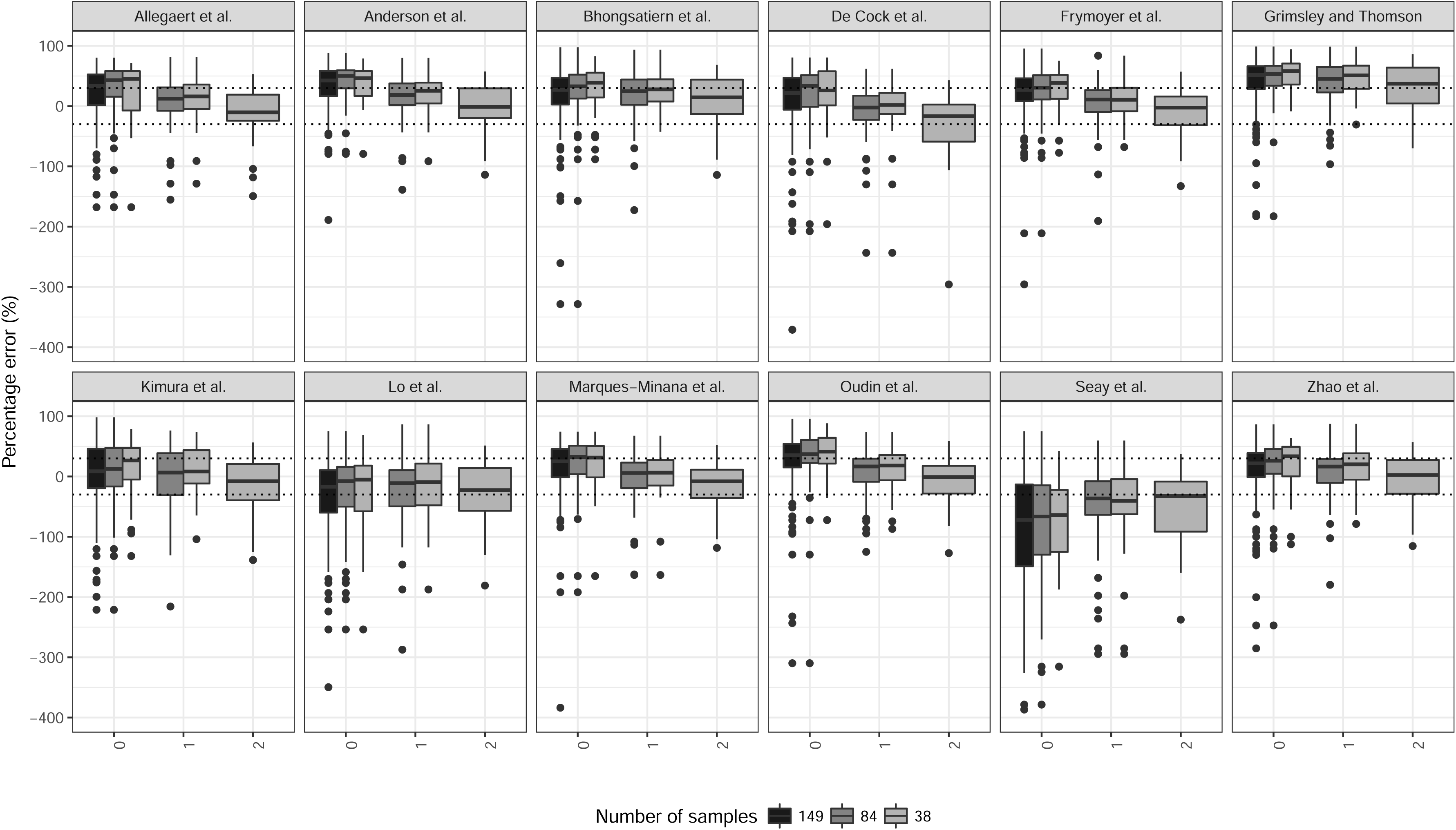
Boxplots of the concentration prediction percentage errors (PE) of 12 vancomycin population PK models. PE distributions are visualized in 3 subsets of individuals. All assayed individuals with at least 1 measured concentration (n = 149) were used to predict the first concentration based on a population model. Subset of individuals with at least 2 concentrations (n = 84) was used to predict the first and second concentrations based on 0 and 1 included individual concentrations, whereas prediction of the third concentration was included for individuals with at least 3 concentrations (n = 38). Dashed lines denote 30% percentage errors

No model attained more than 50% of the PEs within 30% of the true values with no concentration data. The upper limit within 20% was 36.2% with Lo *et al.* [23]. However, predictions including a single concentration obtained more than 50% within 30% in 8 out of 12 models, the highest F30 value was 68% with Marques-Minana *et al.* model [21]. Correspondingly, six models attained more than 40% within 20% of the PEs (Electronic Supplementary Table 3).

### Probability of target attainment simulations

None of the condition-specific error collections used in the adjustment of DosOpt simulated concentrations differed statistically from log-normal distribution (Electronic Supplementary Table 4, Fig. S3). The PTAs for both the 10–15 mg/L and 10–20 mg/L target ranges exceeded the base model estimates (12 models; n = 149) when at least one concentration guided the PK parameter estimation (n = 84). When predictions were made based on two prior concentrations (n = 38), the model-wise PTA estimates remained similar, with attainment probabilities approximately 40% and 60% within 10–15 mg/L and 10–20 mg/L of C_trough_, respectively (Electronic Supplementary Table 5; Fig. 4).

**Fig. 4.**
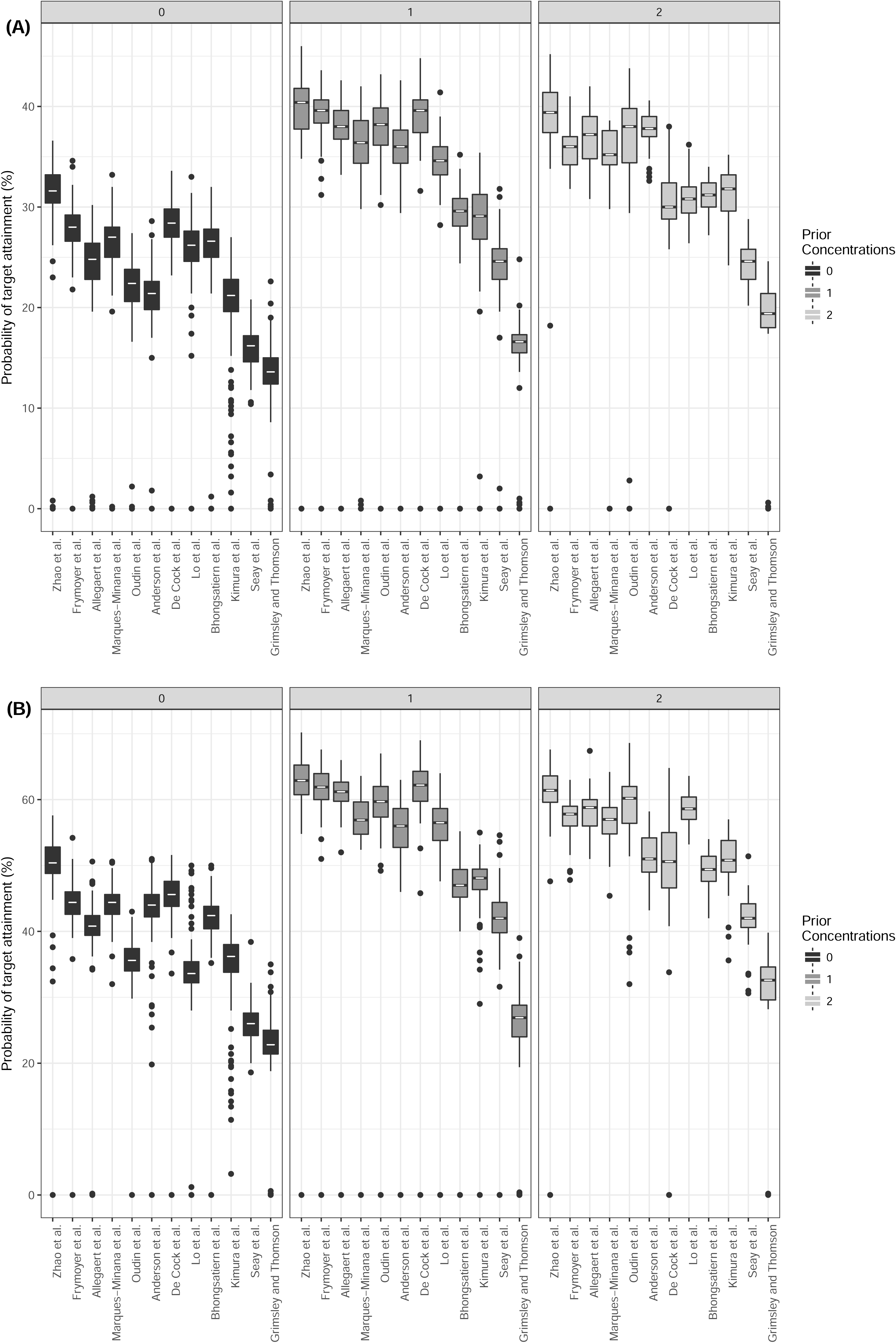
Probability of target attainment estimates with individual pharmacokinetics estimated with 0 (n = 149), 1 (n = 84), and 2 (n = 38) individual concentrations entered in DosOpt. Each point represents the estimated maximal probability of attaining targeted trough concentrations with an individually optimized dose. Simulations assume a dose administered from a baseline concentration of 10 mg/L with a 1-hour infusion and 8-hour interval. Targeted trough concentration ranges (A) 10-20 mg/L, (B) 10-15 mg/L

For example, we observed that a single individual concentration improved the average individual PTAs in the Zhao *et al.* model from 31.6% to 40.4% for target 10–15 mg/L (Fig. 4A) and from 50.4% to 62.9% for target 10–20 mg/L (Fig. 4B). Simulations based on other models also attained relative improvements of around 25% from the population-based predictions. This implies that inclusion of individual concentrations to models with more accurate initial attainment leads to improved achievable PTA.

## Discussion

To the best of our knowledge, this is the first study that compares vancomycin models in neonates whilst also considering individual concentrations in kinetics re-estimation. All published PK models were found biased in population-model based validation but both precision and accuracy improved when including individual concentrations were included. Bayesian inclusion enabled the best performing models to attain around 60% of the concentration predictions within ±30% of the actually measured concentrations. Observed concentration prediction accuracies translated into highly variable simulated target attainment rates, the best performing models achieving around 40% for C*_trough_* 10–15 mg/L and 63% for 10–20 mg/L. These results describe the prediction performances attainable with current models and highlight the importance of model selection for TDM applications like DosOpt. Models underlying precision-based dosing approaches need to be reliably applicable to new populations [32]. Published neonatal vancomycin population PK models have been validated to a different extent. Using an external dataset, we showed that population based NPDE variances were statistically different from expected values in all tested models. Population-based validation of Zhao *et al.* (n = 78) [30] reported similar replication difficulties with only PK models by Grimsley and Thomson and Capparelli *et al.* (did not qualify for this study due to cohort PNA dissimilarity) passing NPDE mean and variance qualification [26,44]. Still, their reported NPDE results were dependent on the assay method used for concentration measurements. This highlights the importance of multiple replications, differences in replication cohorts and complexities of model transferability.

All included models assumed linear PK. Only one [18] was two-compartmental, the rest assumed a single compartment. Uniform model structure limits the feasibility of developing a new population model on our data for component importance assessment. In terms of covariates, all models contained a variable to represent patient age (postnatal, postmenstrual or gestational) and weight (current / birth). The effect of attributes such as creatinine, concomitant medications and artificial ventilation is inconclusive and variably used in models [45]. We hypothesise that continuous dosing used only by Zhao *et al.* and higher similarity of patient age and weights to our retrospective set exemplified by full range cover by Frymoyer *et al.* and Zhao *et al.* [11,20] may have contributed towards improved predictions in our study. However, the small number of evaluated models would leave any attempt at single variable effect estimation statistically under-powered. We also note that two models with consistently the worst predictive accuracy (Seay *et al.* [25], Grimsley and Thomson [26]) were also two of chronologically the oldest. This is a possible indication of improving model development practices.

Large unexplained and inter-individual PK variability of vancomycin in neonates have a negative effect on resulting prediction accuracy. This is illustrated by relatively large residual model component values (Electronic Supplementary Table 1). For example, Zhao *et al.* reported combined error model parameter sizes of 2.28 mg/L for the additive component and 20.3% for the proportional component [20] which add to reported between-subject variability. Individual concentrations help in explaining some uncertainty, but most models still predicted with an absolute error of 20-30% after inclusion of 1 or 2 concentrations. One likely cause is the often un-characterised within-subject PK variability. Another issue would be the failure to incorporate significant covariates in PK models. The extent of such contributory but missing terms is currently unknown as no studies have attempted to partition vancomycin concentration variability into biological and environmental. Nevertheless, identification of novel terms with small effects would require much higher sample sizes than are currently available.

We chose vancomycin C_trough_ ranges of 10-15 mg/L and 10-20 mg/L as the therapeutic targets of PTA simulations as these provide both therapeutic effectiveness and the utility and usefulness in clinical settings. In an experimental study, Ramos-Martin *et al.* suggested that a AUC/MIC higher than 400 is necessary for successful eradication of coagulase negative staphylococci (CoNS)[46]. Padari *et al.* could not correlate AUC/MIC >400 with increased efficacy in neonates treated for CoNS-related bloodstream infections [47]. Studies evaluating the relationship of trough concentration (C_trough_) with AUC have shown that the two measures are not perfect substitutes [7] but relative ease of C_trough_ measurement makes it commonly used for vancomycin therapeutic targeting in guidelines and in clinics [8–10]. Whereas attainment of 10 mg/L C_trough_ has been multiply linked with AUC>400 in paediatric populations [11–15], the estimates on the non-toxic upper end of the PD target ranges are less conclusive and warrant future work. Increased probability of nephrotoxicity has been associated with AUC values exceeding 700 mg*h/L [7,48]. Similarly, some evidence relates C_trough_ >20 mg/L to increased toxicity [49].

Previous assessments of targeted dose adjustment methods have demonstrated the benefit of population PK and Bayesian-based models versus empirical approaches. Compared to the 25.1% of patients achieving 10–20 mg/L C_troughs_ with empirical dosing in a study by Ringelberg [8], Leroux *et al.* reported 72% of patients consistently achieving 15–25 mg/L troughs when the Zhao *et al.* model was used for dose calculations [9]. In a study by Nunn *et al.* [10], Bayesian adjustments resulted in an improvement from 33.8% of patients to 75.0% of patients having a C_trough_ of 10–20 mg/L over the course of a full treatment period. Several *in silico* studies have further showed that Bayesian kinetics estimates improve with the inclusion of additional individual TDM data [10,34,50]. The above studies use a variety of different study designs and C_trough_ ranges to report PTA results, which complicates between-study comparisons. Our simulations similarly showed that inclusion of individual concentrations improved model-wise attainment rates of clinically applicable therapeutic C_trough_ ranges. Inclusion of a single concentration increased the PTA for an average patient from 20–30% to 35–40% in target range 10–15 mg/L (Fig. 3A), and from 35–43% to 55–63% in 10–20 mg/L target range. Firstly, these indicate that development of general Bayesian-guided bedside TDM applications like DosOpt, are beneficial to dose adjustment accuracy. Secondly, the probabilistic attainment of vancomycin therapeutic targets with current models would not provide outcome certainty in the form of PTA values around and above 90% [34].

The present study has several limitations. We did not evaluate concentration performances within patient’s condition severity subgroups, which may have had an unknown effect. The precision of Bayesian parameter estimation may have been influenced by several confounding factors. First, all our samples were assayed using FPIA. Measurement errors and minimal quantifiable concentrations may have varied between our data and in those used for assessed models. Secondly, our samples were overwhelmingly C_trough_. Also, our retrospective dataset consisted of patients with uneven number of available concentrations in comparison groups that complicated comparisons between groups. To assess this effect, we performed additional performance evaluations with 0 and 1 included concentrations for patients with >2 total available concentrations (n = 38), and 0 concentrations for all patients with 2 available concentrations (n = 84).

## Conclusion

Predictions based on previously published vancomycin PK models exhibited considerable performance variability. All population based model predictions were discordant with external validation dataset. However, when at least one treatment concentration was included, both precision and accuracy of predictions improved. Still, relatively low predictive precision of current population models limits attainment of narrow therapeutic targets of vancomycin.

## Notes

### Acknowledgments

This work was supported by institutional research funding grants (IUT34-24) and (IUT34-4). Support from the European Union was also provided through the European Regional Development Fund. We would like to thank Helgi Padari for help with data collection and Hiie Soeorg for constructive comments.

### Conflict of interest

The authors declare no conflict of interests. The authors declare no support from any organization for the submitted work, no financial relationships with any organizations that might have an interest in the submitted work in the previous 3 years, and no other relationships or activities that might have influenced the submitted work.

### Author contributors

TT, TM, and IL designed the study. TT, JV, TM, IL, and RK participated in writing of the manuscript. TT performed the statistical analyses and simulations. TM, RK, and IL contributed and collected the data. JV and IL provided the funding. All authors reviewed the manuscript and approved the final version.

### Compliance with ethical standards

This study was conducted in accordance of good ethical standards. The Ethics Committee of the University of Tartu approved the study in Protocol No. 256/T-23. Ethics committee waived informed consent as the study was retrospectively performed with anonymized data.

## Supplementary Material

Supplementary Material 1 – ESM1.pdf - captions of supplementary tables 1-5 and supplementary figures 1-3

Electronic Supplementary Table 1 - ESMTable1.docx

Electronic Supplementary Table 2 - ESMTable2.xlsx

Electronic Supplementary Table 3 - ESMTable3.xlsx

Electronic Supplementary Table 4 - ESMTable4.xlsx

Electronic Supplementary Table 5 - ESMTable5.xlsx

Fig. S1 - ESMFig1.pdf

Fig. S2 - ESMFig2.pdf

Fig. S3 - ESMFig3.pdf

